# HGFX: a validated Python/JAX reproduction of the Hierarchical Gaussian Filter toolbox

**DOI:** 10.64898/2026.09.22.753415

**Authors:** Mohammad Ahmadkhanloo

## Abstract

The Hierarchical Gaussian Filter (HGF) is a hierarchical Bayesian model of learning under uncertainty and volatility, with a widely used MATLAB implementation. We present HGFX 1.0.0, a Python/JAX toolbox whose primary objective is validated behavioral and numerical compatibility with a frozen HGF Toolbox 8.2.0 reference, while removing MATLAB as a user-runtime dependency. Reimplementation is treated as a validation problem rather than source translation: configuration semantics, parameter transforms, forward trajectories, observation likelihoods, objectives, fitting and statistical surfaces, simulation workflows, official demo behavior, model selection, and backend agreement are compared against the pinned oracle under predeclared tolerances. Two official MATLAB demo workflows reproduce the reference at frozen trajectory tolerances, including a regime in which classic HGF encounters negative posterior precision while eHGF completes successfully. In paired model-selection validation, all 36 BIC winners agree between MATLAB and HGFX. Physical NVIDIA GPU applicability was confirmed on two Tesla T4 devices, with a maximum CPU-versus-GPU final-objective difference of 1.42e-14 against a frozen 1e-7 criterion. Historical parameter-recovery results that did not meet predeclared criteria, together with exact MATLAB/HGFX limitation matches, are preserved and are not reclassified as scientific success. A prospectively gated comparison with pyhgf 0.3.2 shows binary64-scale agreement on mapped perceptual trajectories in one authorized three-level binary-HGF cell, while participant-response negative log-likelihood is retained as not directly comparable. HGFX therefore provides a MATLAB-independent Python implementation with an explicit evidence model that separates direct parity, matched reference limitations, backend applicability, and future performance claims.

## 1 Introduction

Computational models of learning under uncertainty must represent uncertainty about latent states and about the volatility of those states. The Hierarchical Gaussian Filter (HGF) was introduced as a generic hierarchical Bayesian framework for individual learning under uncertainty (C. Mathys et al. 2011) and later developed into a practical filtering framework for perception and learning (C. D. Mathys et al. 2014). It has been used to recover hierarchical prediction errors in neuroimaging (Iglesias et al. 2013) and to model inference about others’ intentions (Diaconescu et al. 2014). The associated MATLAB toolbox is distributed as part of TAPAS, an open-source collection of translational neuromodeling tools (Frässle et al. 2021), and remains a reference implementation for fitting, simulation, model comparison, and analysis.

Reproducing such a toolbox in another language is not equivalent to translating published equations. Scientific behavior also depends on parameter ordering, transforms, prior conventions, fixed and free parameter semantics, placeholder values, numerical primitives, finite-difference behavior, optimizer trajectories, adverse numerical modes, and model-family selection. Small floating-point differences that are negligible at a shared state can be amplified by derivative estimation and non-convex optimization (Wilson and Collins 2019; Peng 2011). A replacement implementation can therefore appear mathematically correct while producing materially different fitted inferences or model-selection outcomes.

Python/JAX toolboxes already exist in this space. pyhgf represents predictive-coding systems as configurable node/edge networks and supports differentiable modern inference (Legrand et al. 2026). HGFX does not claim to be the first Python or JAX HGF. Its v1.0 objective is narrower and complementary: behavioral compatibility with one frozen MATLAB HGF Toolbox 8.2.0 oracle, explicit cross-language evidence, and removal of MATLAB from the user runtime.

This paper asks whether a legacy scientific toolbox can be reproduced in an accelerator-compatible stack without silently changing the scientific contract. We report the released HGFX 1.0.0 evidence: workflow reproduction, paired model-selection agreement, CPU/backend and physical-GPU applicability, a scoped pyhgf comparison, and preserved historical negative results. General speedup and multi-GPU scaling are not headline claims.

For neuroscience methodology, the contribution is a reproducible route for re-running established HGF analyses outside the MATLAB runtime while retaining the validated reference behavior that underpins prior computational-neuroscience and computational-psychiatry applications. The study therefore evaluates software behavior, inferential workflows, and reproducibility rather than introducing a new behavioral or neuroimaging dataset.

## 2 Materials and methods

### 2.1 Software description

HGFX is a Python package (Python >= 3.11) built on NumPy and JAX (Bradbury et al. 2018; Frostig, Johnson, and Leary 2018). The public surface exposes Python-first and MATLAB-style aliases `(fit_model/fitModel, sim_model/simModel, sample_model/sampleModel).` Compatibility-sensitive numerical paths are distinguished from JAX-backed execution. Compatibility repairs were introduced only when supported by an exact MATLAB oracle case and a regression that violated expected behavior.

Software metadata and frozen validation identities are summarized in Table 1.

**Table 1.**
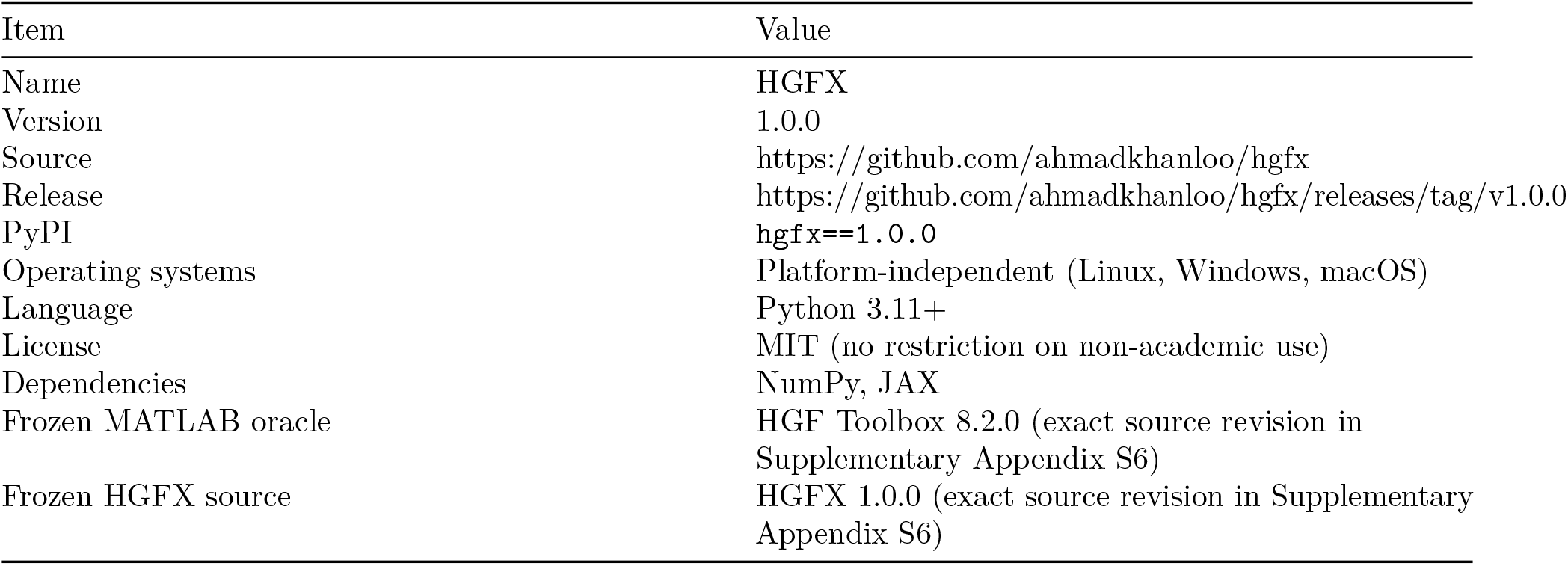
HGFX 1.0.0 software metadata.

Users do not require MATLAB. MATLAB is used only when regenerating cross-language oracle evidence. The HGF Toolbox is cited via Mathys et al. (C. Mathys et al. 2011; C. D. Mathys et al. 2014) and TAPAS (Frässle et al. 2021). Related probabilistic-modelling software includes the VBA toolbox (Daunizeau, Adam, and Rigoux 2014).

### 2.2 Covered HGF families

The HGF represents a hidden hierarchy of Gaussian states in which higher levels encode volatility of lower levels (C. Mathys et al. 2011; C. D. Mathys et al. 2014). For binary observations, a unit-square sigmoid observation model maps the first hidden state to the probability of the observed outcome. HGFX v1.0 covers the classic HGF, the enhanced HGF (eHGF), and the unbounded HGF (uHGF) in the documented MATLAB-compatible configurations, plus specialized surfaces in the v1 migration matrix (sampling, analysis, Bayesian parameter averaging, and the official uHGF-AR(1) demo). Equations are those of the frozen MATLAB toolbox; this article does not introduce a new generative model.

The public fitting interface accepts perceptual and observation configurations, a default transformed start, and a Quasi-Newton optimizer with a frozen iteration budget in the paper protocols. Simulation exports inputs `;u`and responses `y`that become immutable paired inputs when MATLAB is used as an oracle.

### 2.3 Frozen reference and evidence classes

HGFX v1.0 is validated against the frozen HGF Toolbox 8.2.0 reference. Exact source revisions are provided in Supplementary Appendix S6. The compatibility target includes the documented model/configuration inventory, parameter transforms and priors, HGF/eHGF/uHGF and specialized implementations, observation models, fitting and simulation surfaces, Hessian/covariance/correlation/statistical outputs, model selection, official workflow behavior, and public APIs.

We use four interpretation classes:

- **Direct numerical agreement:** MATLAB and HGFX agree under the frozen protocol and tolerance.
- **Matched reference limitation:** the MATLAB oracle exhibits the same numerical or workflow limitation in the same frozen scope; this is not a scientific recovery claim.
- **Preserved negative result:** a predeclared scientific criterion is not met and the result remains archived.
- **Not directly comparable (NDC):** a quantity lacks a defensible common semantic or numerical surface, most relevant to the pyhgf comparison.

Acceptance thresholds, seeds, datasets, starts, model families, optimizers, and grids are not changed after observing results. Results outside predeclared criteria remain archived. The numerical policy and the two most sensitive optimizer/basin cases are documented in Supplementary Appendices S1-S2.

### 2.4 Validation hierarchy

The evidence hierarchy is: frozen reference identity; machine-readable validation artifacts; paired MATLAB/HGFX raw outputs; continuous-integration provenance; and manuscript tables/figures generated from those artifacts. Numerical checks use IEEE-754 binary64. The project does not target arbitrary-precision solutions that differ from frozen MATLAB execution.

Two official MATLAB demo workflows (320 binary trials) are release-gated. Complete release-relevant trajectories and inference states are compared at `rtol = 5e-11` and `atol = 5e-13`.

Fitting validation covers objectives at fixed parameters, MATLAB-compatible optimizer behavior, fitted parameters where direct parity is expected, trajectories, predictions/residuals, Hessian-derived covariance/correlation, and AIC/BIC/LME.

An earlier frozen parameter-recovery experiment did not meet its scientific acceptance criteria and remains preserved. Subsequent paired validation distinguishes parameter recovery from model selection on a frozen three-model grid (three binary perceptual models; truth scales 0.15 and 0.35; Quasi-Newton; predeclared thresholds: convergence rate >= 0.80, median correlation >= 0.50, median standardized RMSE < = 1.00, model-recovery balanced accuracy >= 0.50).

CPU/backend equivalence and physical NVIDIA GPU applicability were tested after an independent review of portability blockers (host C runtime `expm1`/`log` paths and Windows CRLF hash mismatches). Both blockers were remediated without changing scientific thresholds.

### 2.5 pyhgf common-scope protocol

`pyhgf==0.3.2` was pinned before execution. Direct comparison was allowed only after model structure, update equations, parameters, initialization, input/masking semantics, reported quantities, precision mode, and numerical guards had been mapped. The authorized cell is a fixed-parameter, fully observed, three-level binary HGF (128 trials; standard/mean-field updates; unit coupling; zero drift; inverse temperature `ze = 48`). Trajectory quantities used `atol = 1e-10` and `rtol = 1e-8`; total response NLL used `atol = 1e-7` and `rtol = 1e-8`. Raw arrays were hashed before interpretation. A derived response-NLL boundary nonfinite was prospectively classified as NDC rather than triggering post-result clipping.

### 2.6 Prospective trial-horizon protocol

A prospectively frozen paired MATLAB/HGFX trial-horizon experiment examined 128, 256, 512 and 1024 trials using the same three perceptual models, truth scales, seeds, Quasi-Newton budget, and predeclared recovery thresholds. The preregistered scientific comparison was 256 versus 1024 after a paired-integrity gate; 128 and 512 were trajectory diagnostics. Invalid simulations and fits outside the preregistered criteria were retained rather than resampled. The experiment completed all 24 preregistered shards. The paired-integrity requirement was not satisfied: 10 of 72 model-recovery BIC winners disagreed, all for the classic binary HGF at 512 or 1024 trials, and parameter metrics for those horizons were undefined because invalid simulations were retained. The evidence was therefore judged insufficient to support a data-horizon or structural-identifiability conclusion.

## 3 Results

### 3.1 Workflow equivalence

HGFX 1.0.0 was used for all release-level comparisons. Official demo reproductions execute the MATLAB oracle and the Python reproduction on matched inputs, compare complete release-relevant outputs, and archive machine-readable artifacts (Table 2). Exact source identity is reported in Supplementary Appendix S6.

**Table 2.** Official workflow and analysis-surface coverage. Machine-readable source data are available in the reproducibility package.

| Surface | Classification |
| --- | --- |
| Core model/API compatibility (documented scopes) | Agreement within documented compatibility scope |
| uHGF to AR(1) official workflow | Reference trajectory reproduced within frozen tolerance |
| sampleModel / prior-predictive workflow | Reference workflow reproduced |
| Correlation/residual analysis surfaces | Reference outputs reproduced within frozen tolerances |
| Bayesian parameter averaging | Reference outputs reproduced within frozen tolerances |

The first demo is a regime in which classic binary HGF encounters negative posterior precision while eHGF succeeds. HGFX reproduces this: `hgf_binary` reaches negative posterior precision in both implementations, `ehgf_binary` completes in both, and eHGF trajectories agree within the frozen tolerance (workflow parity).

The second demo reproduces the uHGF to uHGF-AR(1) transition. The maximum absolute third-level posterior mean is 16.99162398501939 for uHGF and 4.0927117005012175 for uHGF-AR(1) in both implementations (direct uHGF-AR(1) workflow parity).

### 3.2 Fitting statistics and scoped limitations

Two fitting-validation cases expose reference limitations rather than direct fitting parity (Table 3). In an official fitting stress case, MATLAB and HGFX can terminate at different optimizer endpoints in a numerically sensitive basin. In a separate frozen Level-2 holdout case, the inference-equivalence criterion is not met for a specific seed. In both cases the frozen MATLAB oracle shows the corresponding instability or sensitivity, so these results are reported as matched reference limitations rather than as successful parameter recovery.

**Table 3.** Fitting and recovery classifications. Direct mismatches and negative results are retained.

| Surface | Direct result | Paper disposition |
| --- | --- | --- |
| Earlier frozen parameter-recovery experiment | scientific criteria not met | preserved negative result |
| Official fitting stress case | optimizer endpoint mismatch in sensitive basin | MATLAB reference shows the same start sensitivity |
| Frozen Level-2 holdout fit | inference-equivalence criterion not met for a fixed seed | MATLAB reference shows the same start sensitivity |
| Paired parameter recovery | recovery criteria not fully met in either implementation | paired numerical behavior closely reproduced |
| Paired model selection | 36/36 BIC winners match | direct agreement |

### 3.3 Parameter recovery versus model selection

Paired parameter-recovery summaries are shown in Figure 1 and Table 4. MATLAB and HGFX closely reproduce the same recovery summaries, but the predeclared scientific recovery criteria are not fully met. Paired model selection is stronger: all 36/36 BIC winner decisions match between MATLAB and HGFX in the same frozen grid. Model-selection agreement is not used to imply strong parameter identifiability. Full grid dimensions, frozen thresholds, and the prospective trial-horizon extension are reported in Supplementary Appendix S3.

**Table 4.** Paired parameter recovery on the frozen three-model grid (both truth scales). Thresholds: convergence >= 0.80, median correlation >= 0.50, median standardized RMSE <= 1.00. Full-precision values are available in the reproducibility package.

| Model | MATLAB / HGFX<br>convergence | MATLAB / HGFX<br>median r | MATLAB / HGFX<br>median sRMSE | Criteria outside<br>target |
| --- | --- | --- | --- | --- |
| hgf_binary | 0.833 / 0.833 | 0.203 / 0.203 | 2.610 / 2.610 | correlation, sRMSE |
| ehgf_binary | 0.917 / 0.917 | 0.462 / 0.462 | 2.359 / 2.359 | correlation, sRMSE |
| uhgf_binary | 0.792 / 0.792 | 0.381 / 0.381 | 2.958 / 2.958 | convergence,<br>correlation, sRMSE |

**Figure 1:**
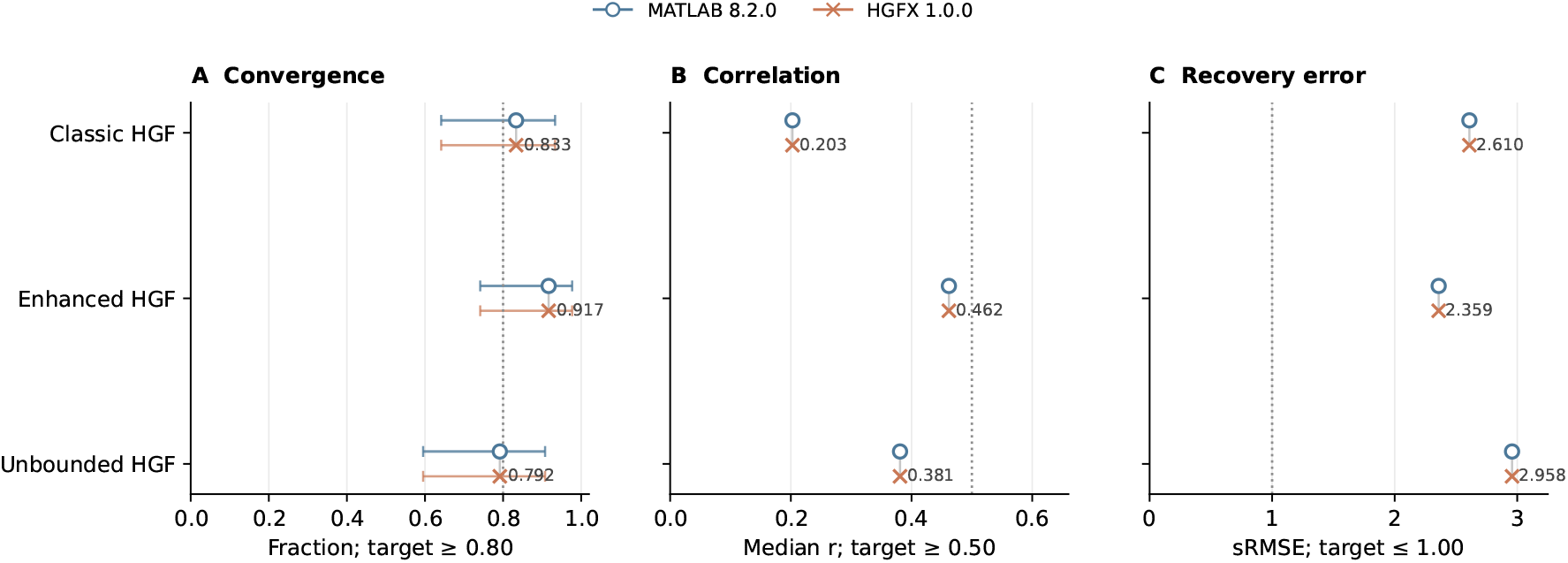
Close paired agreement coexists with weak recovery. Original-scale parameter-recovery summaries are shown for classic, enhanced, and unbounded HGF. Circles denote MATLAB 8.2.0 and crosses denote HGFX 1.0.0; dotted lines mark the unchanged targets. The convergence panel includes descriptive Wilson 95% intervals for 24 cases per family. Correlation and standardized-RMSE panels show point summaries because replicate-level bootstrap inputs are not available in the archived aggregate.

The trial-horizon study (section 2.6) is complete as an executed protocol, not as a positive identifiability result. Because its paired-integrity requirement was not satisfied, the available evidence is insufficient for a stronger recovery or identifiability conclusion. Diagnostic per-horizon values are shown in Figure 2; they do not replace the earlier recovery evidence.

**Figure 2:**
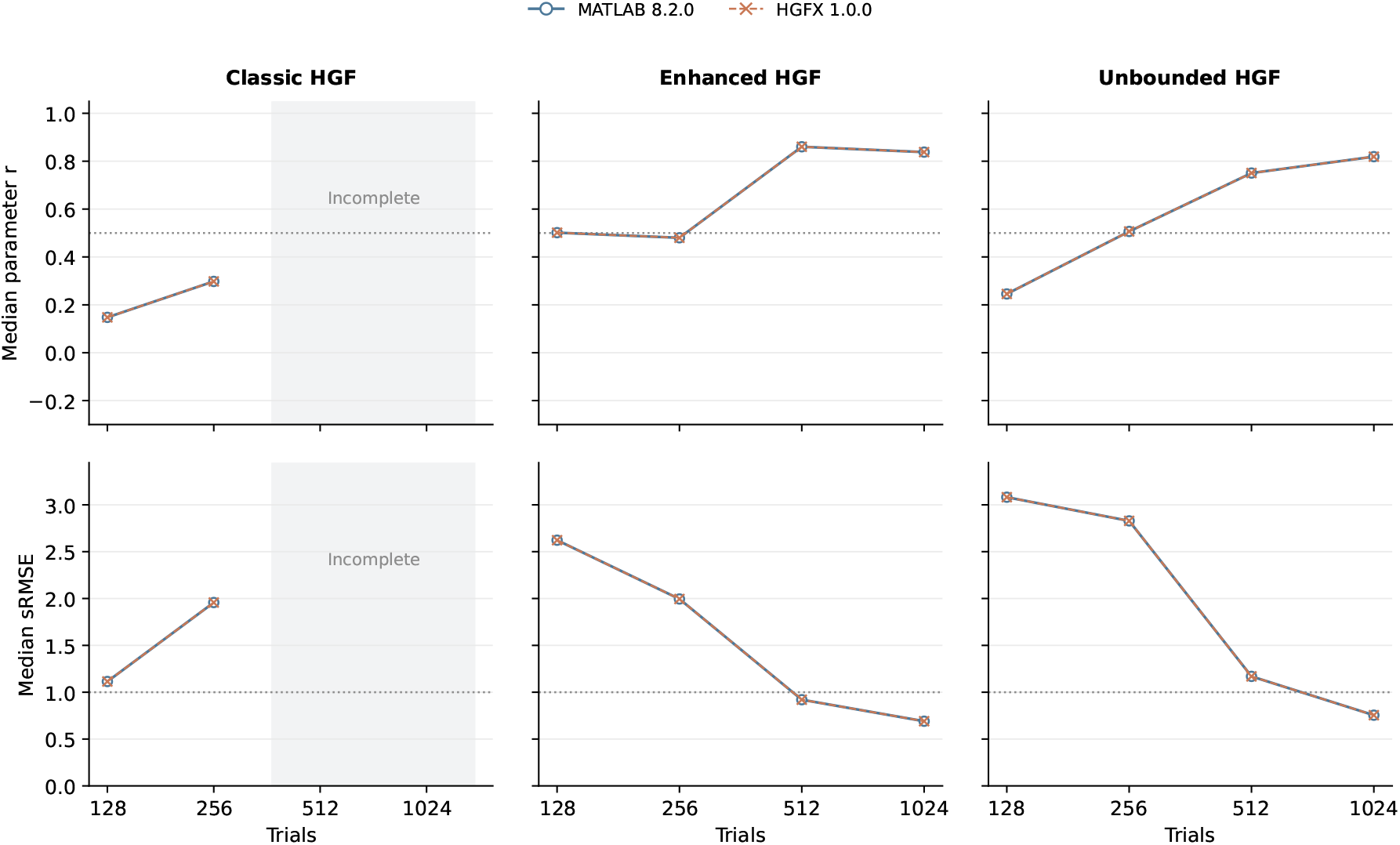
Trial-horizon diagnostics with incomplete cells retained. Columns separate classic, enhanced, and unbounded HGF; rows show median parameter correlation and median standardized RMSE at 128, 256, 512, and 1024 trials. Dotted lines mark the original targets. Shaded regions denote incomplete classic-HGF summaries at 512 and 1024 trials and are not zeros or interpolated estimates. The whole-study paired-integrity requirement was not satisfied, so these trends do not establish identifiability.

### 3.4 Backend and physical-GPU applicability

Compatibility CPU and JAX-backed CPU outputs agree under the frozen backend-equivalence criterion. On two Tesla T4 GPUs (Python 3.12.13, JAX/JAXLIB 0.11.1, `nvidia-smi` process residency), all four required CPU-versus-GPU fitting cells remain below the predeclared final-objective difference criterion of 1e-7. The maximum gap is 1.4210854715202004e-14 (Table 5). This is applicability/correctness evidence, not a speed or scaling result. Exact hardware/runtime provenance, the four CPU-versus-GPU objective pairs, the execution command, and the raw-artifact checksum are reported in Supplementary Appendix S5.

**Table 5.** Backend and GPU applicability.

| Surface | Classification | Result |
| --- | --- | --- |
| Compatibility vs JAX CPU | direct backend agreement | objective/backend agreement |
| Physical NVIDIA GPU (2x Tesla T4) | physical-GPU applicability confirmed | max abs objective gap = 1.42e-14; criterion 1e-7 |

### 3.5 Common-scope comparison with pyhgf

In the authorized 128-trial cell, all 11 mapped perceptual/inference quantities were within predeclared tolerances (Figure 3). Maximum absolute differences were at binary64 rounding scale (1.11e-16 to 1.55e-15). Observed-input integrity was exact.

**Figure 3:**
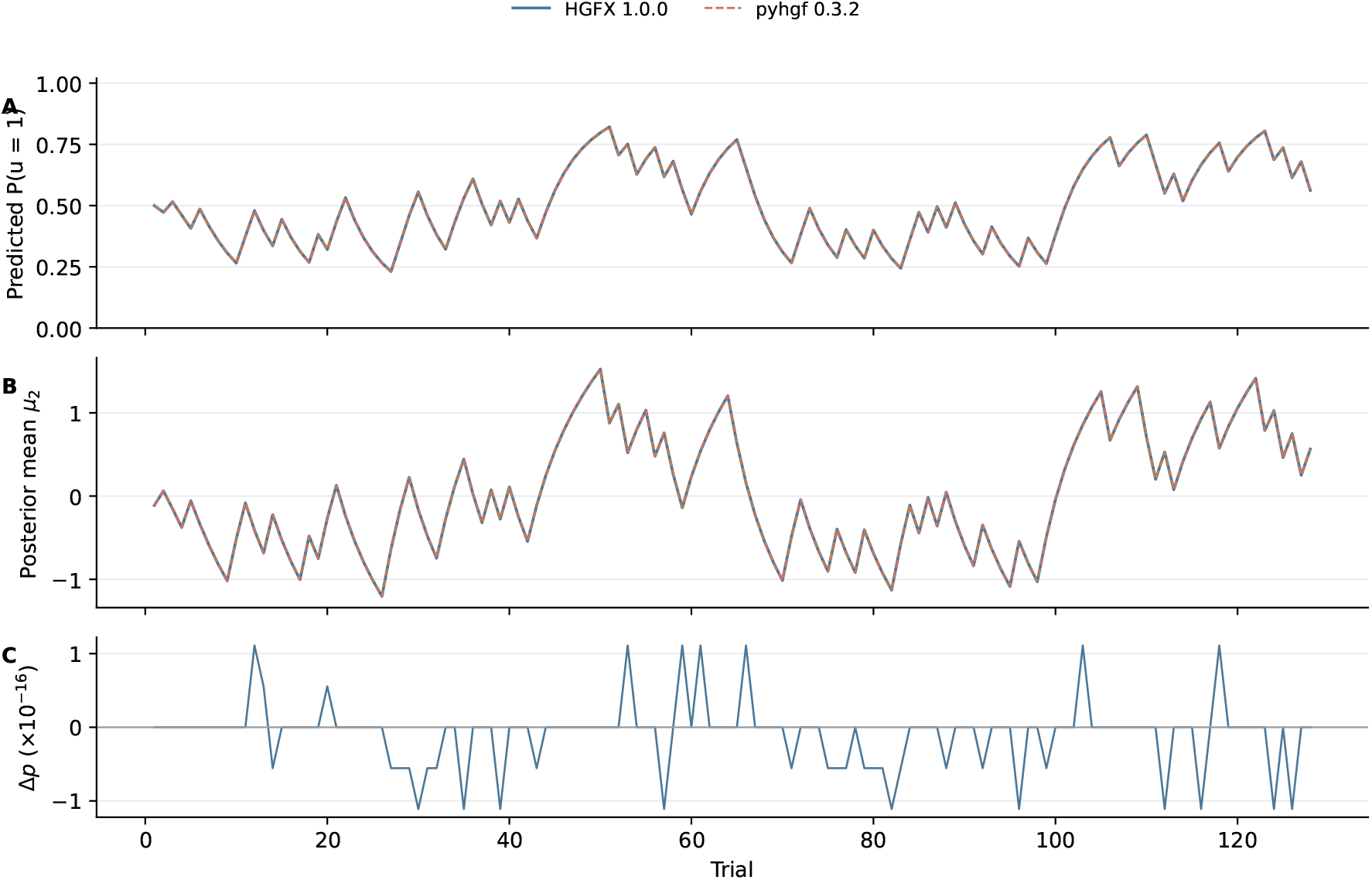
Perceptual trajectories agree at binary64 rounding scale, with residuals shown separately. HGFX 1.0.0 and pyhgf 0.3.2 are compared on the frozen 128-trial three-level binary-HGF cell. Panels show predicted input probability, level-2 posterior mean, and the signed HGFX-minus-pyhgf probability residual in units of 1e-16. Participant-response NLL is a separate quantity and remains not directly comparable in this cell.

Participant-response NLL was not directly comparable. With `ze = 48`, the pyhgf-side power-ratio transformation reached an exact probability boundary on 13 trials and unclipped surprise became +Inf; the HGFX log-domain `unitsq_sgm` evaluation remained finite (total NLL 1808.855415351429). No clipping, formula, precision, parameter, input, or tolerance was changed after observing the result. Overall interpretation: mapped perceptual quantities agree in the authorized cell, while participant-response NLL remains not directly comparable. The semantic gate, exact comparator identity, and numerical-boundary details are given in Supplementary Appendix S4.

### 3.6 Examples of use and current limitations

Typical use after `pip install hgfx==1.0.0` is MATLAB-style fitting and simulation from Python, including the official demo reproductions. MATLAB is unnecessary for those user paths. Current limitations: (i) compatibility is scoped to HGF Toolbox 8.2.0, not future upstream versions; (ii) two numerically sensitive fitting cases are matched reference limitations rather than scientific recovery successes; (iii) parameter identifiability is weaker than paired model selection, and the trial-horizon experiment did not satisfy its paired-integrity requirement, so it does not support a stronger recovery claim; (iv) GPU evidence is T4 applicability, not throughput; and (v) the pyhgf result is one mapped cell, not package-wide equivalence.

## 4 Discussion

Reproducing a scientific toolbox requires a broader notion of compatibility than implementing published equations. In numerically sensitive fitting problems, binary64-scale elementary differences can be amplified through finite-difference derivatives and quasi-Newton optimization, producing different endpoints even when shared-state objectives are extremely close.

This motivates separating scientific correctness from reference faithfulness. When the MATLAB oracle is itself unstable in a tested scope, forcing Python toward a preferred endpoint can be less faithful than preserving the oracle’s limitation. A matched limitation is not evidence that the recovered parameter is identifiable. HGFX therefore preserves the original negative results and reports the narrow product-compatibility interpretation separately.

Workflow-level validation matters for the same reason. Reproducing the expected negative-posterior-precision behavior of classic HGF in a documented regime, while reproducing successful eHGF behavior, is part of compatibility. Treating every adverse or unstable outcome as an implementation bug would have encouraged divergence from the reference.

pyhgf and HGFX overlap but have different design centers. pyhgf emphasizes generalized network construction and differentiability (Legrand et al. 2026). HGFX v1.0 emphasizes frozen-MATLAB compatibility and provenance. Quantity-specific claims are more informative than ranking the packages. Mapped belief trajectories agree to rounding scale in the authorized cell; the response-NLL surface exposes a numerical-boundary difference despite sharing the same predicted belief.

JAX enables compiled CPU/GPU execution, batching, and multi-device workloads (Bradbury et al. 2018; Frostig, Johnson, and Leary 2018). Accelerator-native code is scientifically useful only if the accelerated path preserves relevant outputs. The T4 result supports that claim for the tested objective. Throughput depends on workload size, compilation amortization, hardware, device count, and contention; protocol 1 therefore does not activate a headline performance or scaling result. Scalability of the JAX path is an engineering capability of the implementation, not a reported empirical law.

The strongest contribution is methodological: an explicit oracle, evidence classes, a numerical compatibility policy, preservation of negative results, and release provenance. That discipline reduces the risk that a modern implementation gains convenience by silently changing an established scientific contract.

## 5 Data and code availability

HGFX 1.0.0 is MIT-licensed (HGFX developers 2026). Source, tag, and package URLs are in Table 1. All tables and figures are generated from committed machine-readable evidence by scripts included with the source repository. MATLAB is required only to regenerate paired oracle evidence. Exact source revisions, repository-facing case identifiers, checksums, workflow provenance, and claim-to-evidence traceability are intentionally confined to Supplementary Appendix S6 and the reproducibility package.

## Supplementary material guide

The supplementary appendices are intentionally separated from the main narrative so that technical audit detail does not obscure the scientific results. They are organized as follows:

- **S1 — Numerical compatibility policy:** frozen tolerances, numerical rules, and acceptance logic.
- **S2 — Numerically sensitive fitting cases:** detailed evidence for the two optimizer/basin-sensitive workflows.
- **S3 — Recovery and model-selection protocol:** full grid design, thresholds, parameter-recovery results, paired model-selection results, and the prospective trial-horizon extension.
- **S4 — pyhgf common-scope comparison:** semantic mapping, numerical comparison, and the response-NLL non-comparability boundary.
- **S5 — Physical-GPU applicability:** hardware/runtime provenance, CPU-versus-GPU objective pairs, execution command, and limitations.
- **S6 — Reproducibility and traceability:** exact source revisions, internal case identifiers, workflow provenance, hashes, and links from reader-facing claims to repository evidence.

Each appendix is cited at the point where its detail becomes relevant; S6 is the audit trail rather than part of the scientific narrative.

## CRediT authorship contribution statement

Mohammad Ahmadkhanloo: Conceptualization, Software, Validation, Formal analysis, Data curation, Visualization, Writing – original draft, Writing – review & editing.

## Acknowledgments

The frozen MATLAB HGF Toolbox 8.2.0 of Mathys and colleagues is the reference oracle for this work. GitHub Actions and MATLAB were used only to regenerate paired oracle evidence.

## Supplementary Appendices

These appendices accompany **HGFX: a validated Python/JAX reproduction of the Hierarchical Gaussian Filter toolbox**. They provide the numerical-policy, sensitivity, provenance, and traceability details that are intentionally summarized in the main text. Repository identifiers are confined to the final traceability appendix so the scientific narrative remains reader-facing.

### Appendix S1. Numerical compatibility policy and frozen acceptance rules

HGFX treats MATLAB compatibility as a numerical validation problem rather than a source-translation exercise. The frozen reference is HGF Toolbox 8.2.0 at commit `2437f4dc241541072722a2695ddeca7b44d83dd3`; the released HGFX source is v1.0.0 at `4dd8fbd8239d05f2c7932a9a9b3b7795f0a9ab27`.

All paper-facing acceptance thresholds, seeds, datasets, model families, starts, optimizers, and validation grids were fixed before interpretation of the corresponding final results. Negative, failed, and not-directly-comparable outcomes were retained. No failed replicate was silently dropped or regenerated, and no tolerance was widened after observing results.

Official release-relevant trajectory comparisons use IEEE-754 binary64 with `rtol = 5e-11` and `atol = 5e-13`. The prospective recovery criteria are unchanged from the frozen protocol:

- parameter convergence rate >= 0.80;
- median finite parameter correlation >= 0.50;
- median standardized RMSE <= 1.00;
- model-recovery balanced accuracy >= 0.50.

The physical-GPU applicability criterion is a CPU-versus-GPU final-objective absolute gap <= `1e-7`.

The paper distinguishes four reader-facing interpretations: direct parity, matched reference limitation, preserved failure, and not directly comparable. A matched reference limitation is never promoted to parameter-recovery success or direct fit parity.

### Appendix S2. Numerically sensitive fitting cases

#### S2.1 Official enhanced-HGF fitting stress case

In one official enhanced-HGF fitting workflow, MATLAB and HGFX can terminate in different optimizer basins even though shared-state numerical agreement remains at binary64 scale.

The exact shared-vector objective checks pass under the unchanged numerical gate. At the first localized off-centre sample, the objective absolute difference is `2.842170943040401e-14`, localized to the observation log-likelihood. The first-trial log-likelihood absolute difference is`2.220446049250313e-16`,the maximum per-trial log-likelihood absolute difference is 8.881784197001252e-16, and the MATLAB vector-sum versus scalar-loop reduction difference is `1.1368683772161603e-13`. The inference-state maximum absolute difference at the localized source-likelihood probe is exactly zero.

A MATLAB self-sensitivity probe perturbed the official transformed start by one local floating-point spacing. All 6 independent +/-one-spacing perturbations produced materially different optimizer endpoints, despite preserving the same model, data, optimizer, and scientific tolerances. The largest observed free-parameter shift was approximately 1.508. This demonstrates that the residual endpoint mismatch is consistent with a numerical-basin sensitivity also present in the frozen MATLAB reference; it is therefore reported as a matched reference limitation, not as inferential parity.

#### S2.2 Frozen uHGF Level-2 holdout case

A separate uHGF + Gaussian-observation holdout used a preselected seed and unchanged Level-2 tolerances `rtol = 3e-8` and `atol = 3e-10`. The frozen holdout fails inference equivalence for that seed, while exact shared-state objective/gradient checks and exact-state quasi-Newton/BFGS replay do not identify a material HGFX-only semantic defect.

The MATLAB reference is itself strongly start-sensitive in this exact workflow: 13 of 14 independent +/-one-local-spacing perturbations move the MATLAB optimizer endpoint outside the unchanged endpoint gate. The failed holdout therefore remains visible as a failure and is interpreted only as an exact-scope matched reference limitation. The result does not justify changing the seed, start, optimizer, data, or tolerance.

### Appendix S3. Recovery and model-selection protocol

The paired recovery program separates parameter recovery from model recovery.

For the frozen paired recovery grid used in the main paper, the perceptual models are classic binary HGF, enhanced binary HGF, and unbounded binary HGF, with the `unitsq_sgm` observation model. The grid contains 72 parameter-recovery cases and 36 model-recovery datasets; each model-recovery dataset is fit by all three candidate models. Trial counts are 128 and 256, truth perturbation scales are 0.15 and 0.35 prior standard deviations, parameter recovery uses 6 replicates per stratum, and model recovery uses 3 replicates per stratum.

The resulting parameter-recovery summary is intentionally not labeled a scientific PASS. MATLAB and HGFX agree closely on the frozen metrics:

| Model | Convergence<br>MATLAB/HGFX | Median r<br>MATLAB/HGFX | Median sRMSE<br>MATLAB/HGFX |
| --- | --- | --- | --- |
| classic binary HGF | 0.833 / 0.833 | 0.203 / 0.203 | 2.610 / 2.610 |
| enhanced binary HGF | 0.917 / 0.917 | 0.462 / 0.462 | 2.359 / 2.359 |
| unbounded binary HGF | 0.792 / 0.792 | 0.381 / 0.381 | 2.958 / 2.958 |

By contrast, paired model selection is stronger: all 36 of 36 BIC winners agree between MATLAB and HGFX, and balanced accuracy is 0.5833333333333334 in both implementations. This agreement is not used to imply strong parameter identifiability.

#### S3.1 Prospective trial-horizon extension

A separately frozen prospective extension examined 128, 256, 512, and 1024 trials, with the same three perceptual models and truth scales. Parameter recovery used 6 replicates per model x horizon x truth-scale stratum. Model recovery used 3 replicates per generating-model x horizon x truth-scale stratum. The frozen workload was 360 fits per implementation.

The paired-integrity gate required complete paired shards, identical free-parameter indices, identical pass/fail outcomes for each frozen parameter criterion at each model x horizon, and agreement of every paired BIC winner before any identifiability interpretation.

That gate did not pass. Ten of 72 BIC winners disagreed, all in classic binary-HGF datasets at 512 or 1024 trials. Parameter-recovery summaries for classic HGF at those horizons are incomplete because invalid simulations were retained rather than resampled. Consequently, the study supports no data-horizon or structural-identifiability conclusion. This negative outcome is retained rather than repaired post hoc.

### Appendix S4. pyhgf common-scope semantic and numerical comparison

The external comparator was frozen as `pyhgf==0.3.2` before execution, with source-distribution SHA-256

`8289f6746668e3af9878c3b5638484c70cd44a596e796ec281986da47e9c723d`.

Direct comparison was authorized only after mapping model structure, update equations, input/masking semantics, parameter meanings and transforms, initial states, observation/response quantities, precision mode, and numerical guards. The executed cell is a fully observed fixed-parameter three-level binary HGF with 128 trials and `ze = 48`.

All 11 mapped perceptual/inference quantities pass their prospectively frozen tolerances. Maximum absolute errors span approximately `1.11e-16` to `1.55e-15`, i.e. binary64 rounding scale. Observed inputs match exactly.

Participant-response negative log-likelihood is not directly comparable in this cell. The pyhgf-side power-ratio response transformation reaches an exact probability boundary on 13 trials, yielding nonfinite un-clipped surprise, whereas the HGFX log-domain `unitsq_sgm` evaluation remains finite with total NLL `1808.855415351429`. No clipping, formula, precision, parameter, input, version, or tolerance was changed after observing the result.

This is a quantity-specific comparison, not a package ranking and not evidence of package-wide equivalence.

### Appendix S5. Physical-GPU applicability and environment provenance

Physical GPU validation was executed in a hosted/shared Kaggle environment with two NVIDIA Tesla T4 devices. The recorded runtime was Python 3.12.13, JAX 0.11.1, and JAXLIB 0.11.1. Physical CUDA residency was verified through the runtime device list and `nvidia-smi` process evidence.

Four required CPU-versus-GPU fitting cells were evaluated:

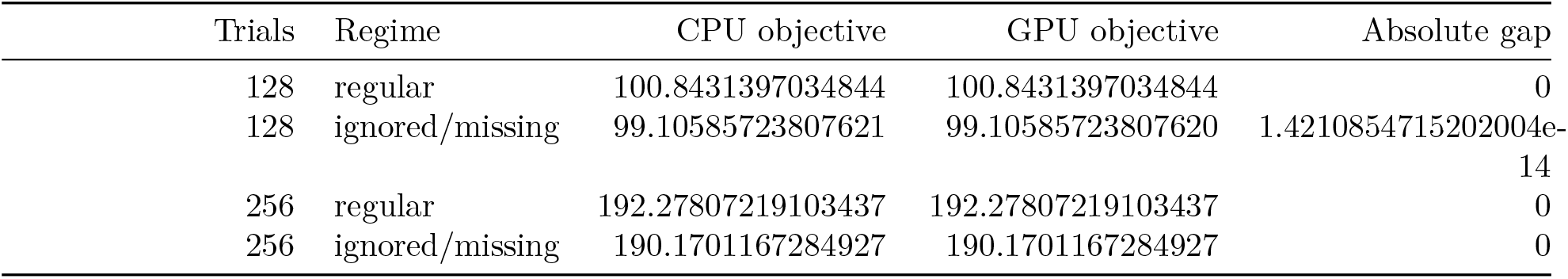

|  | Trials | Regime | CPU objective | GPU objective | Absolute gap |
| --- | --- | --- | --- | --- | --- |
|  | 128 | regular | 100.8431397034844 | 100.8431397034844 | 0 |
|  | 128 | ignored/missing | 99.10585723807621 | 99.10585723807620 | 1.4210854715202004e-14 |
|  | 256 | regular | 192.27807219103437 | 192.27807219103437 | 0 |
|  | 256 | ignored/missing | 190.1701167284927 | 190.1701167284927 | 0 |

All four are below the frozen `1e-7` final-objective criterion. The maximum observed gap is 1.4210854715202004e-14.

This evidence establishes physical-GPU applicability/correctness only. The environment was shared, and no claim of uncontended peak performance, general speedup, H100 performance, or multi-GPU scaling is made.

The exact recorded command was:

`/usr/bin/python3 /kaggle/working/hgfx/scripts/run_m18_s9_backend_robustness.py --output`

The preserved raw validation artifact SHA-256 is `6cd35c82be1e542830c06f6b7b7e444fda0ff4fe93773f080e6c13725212dfdf`

### Appendix S6. Reproducibility and repository traceability

The main text deliberately avoids internal milestone/case identifiers. They are listed here only to provide an auditable bridge from reader-facing claims to repository evidence.

| Reader-facing item | Repository identifier / evidence |
| --- | --- |
| Official enhanced-HGF fitting stress case | D02;<br>reference/validation/m18_d02_reference_limitation/decision |
| Frozen uHGF Level-2 holdout case | D08;<br>reference/validation/m18_d08_reference_limitation/decision |
| Paired recovery/model-selection grid | S7;<br>reference/validation/m18_s7_reference_limitation/decision |
| Physical GPU applicability | S9;<br>gpu_validation_results/m18_s9_physical_gpu_revalidation.js |
| Prospective trial-horizon extension | M18C.2/P3;<br>paper/reproducibility/p3_m18c2_aggregate_35272347167.json |
| pyhgf common-scope comparison | P2A.10;<br>paper/reproducibility/p2a10_comparison_35268575414.json |

Primary reproducibility identities:

1. HGFX release: `v1.0.0` / `4dd8fbd8239d05f2c7932a9a9b3b7795f0a9ab27`;
2. MATLAB oracle: HGF Toolbox 8.2.0 / `2437f4dc241541072722a2695ddeca7b44d83dd3`;
3. paper protocol: `hgfx-paper-protocol-1`;
4. trial-horizon Actions run: `35272347167`;
5. trial-horizon aggregate SHA-256: `83ccbb7f5c4f0eed213d60330d0b318a37e74f03ba08a93a4e4d5d60841131b4`;
6. pyhgf comparison raw-result SHA-256: `202007865c78ba0b138eeda5f105a73399d74076a802edd2c422ddcf98e4696b`.

Paper tables and figures are generated from committed machine-readable evidence by committed scripts. The reviewer entry point is `paper/reproducibility/README.md`. MATLAB is required only to regenerate paired oracle evidence; it is not required to install or use HGFX.

